# Global scientific research in Microbial Induced Calcite Precipitation: A bibliometric network analysis

**DOI:** 10.1101/2022.10.13.512087

**Authors:** Mazhar Ali Jarwar, Stefano Dumontet, Rosa Anna Nastro, M. Esther Sanz-Montero, Vincenzo Pasquale

## Abstract

Microbial Induced Calcite Precipitation (MICP) offers a host of interesting features, both from theoretical and practical standpoints. This process was firstly investigated as a geo-biological mechanism involved in carbonate mineral formation both in rocks and soil. The interest towards its practical use significantly increased in the recent years, as MICP was used in different fields, such as oil recovery, improvement of soil geotechnical characteristics and concreate healing. To the best of our knowledge, this paper is the first attempt to carry out a bibliometric descriptive study of publications on MICP. We analyzed data from the Web of Science Core Collection (WoSCC), which provides comprehensive information for bibliometric analysis, including the Science Citation Index-Expanded (SCI-E) and the Social Sciences Citation Index (SSCI). The bibliometric analysis was carried out on 1580 publications, from 2000 to August 2022, and included publication output, author, institution, country, collaborations between authors, institutes and countries, and citation frequency. We created visualization maps including research collaborations by using the VOSviewer program. MICP, carbonate precipitation, cementation, and soil improvement in terms of geotechnical properties are high-frequency used keywords. Although in the year 2000 only two papers were published on MICP, the number of publications increased rapidly since 2014. In 2021, 333 papers were published. China leads as the most productive country, followed by USA and Australia. According to our results, the number of research papers dramatically increased in the last 5 years. MICP with concrete healing/cementation, soil geotechnical improvement, and the low environmental impact of such a technique are becoming very popular topics among researchers. With the ageing of concrete buildings, as well as with the worsening of environmental pollution and soil alterations, the research about MICP will play an ever increasing and crucial role in civil engineering and geotechnical issues and in soil science. Nevertheless, our study pointed out a sort of concentration of the MICP studies in few countries. Russia and Brazil, for instance, seems to poorly contribute to MICP researches. A larger cooperation among countries, together with the extension of the research network on this topic, would considerably foster the progress in MICP studies, from both practical and speculative standpoints.

## 1. Introduction

Carbonates represent a major reservoir of carbon on the Earth (about 41.9% of the total carbon on our planet) and Microbially Induced Calcite Precipitation (MICP) plays a prominent role within the global carbon cycle (Zhu and Dittrich, 2016). The study of the role of microbial mediated mineral precipitation in natural environment takes place at the interface between microbiology and geology and includes the understanding of both autotrophic and heterotrophic metabolic pathways. MICP played a crucial role in mineral deposition during geological eras, as proved by geological record (Dupraz et al., 2009; Laméland et al., 2022; Hoffmann et al., 2021). Calcium carbonate minerals forming stalactites, stalagmites, microbialites, stromatolites and thrombolites represent examples of minerals produced by the activities of microorganisms and the overall effect of such a microbial activity extends up to landscapes morphology (Hoffman et al., 2021; Dupraz et al, 2009).

The first published paper dealing with the ability of bacteria to induce carbonate precipitation dates back to the early 19^th^ century (Görgen et al., 2021). Later, the ability to precipitate CaCO_3_ has been considered as general phenotype among bacteria (Boquet et al., 1973; Krumbein 1974). According to Dupraz et al. (2009), MICP includes two different processes: microbially-induced mineralization (where metabolic activities of living microorganisms induce conditions for mineral precipitation) and biologically-influenced mineralization (where dead or metabolically inactive microbial cells, together with environmental factors, lead to a passive precipitation of minerals).

The precipitation of Ca carbonates and, more in general of Ca minerals, is shared among different domains of microorganisms other than bacteria. Cyanobacteria (Kamennaya et al., 2012; Lamérand et al., 2022; Görgen et al., 2021), fungi (Pasquale et al., 2019), algae (Nash et al., 2019; Samuels and al., 2020; Mulec et al., 2008), and even some vascular plants (Ostrofsky and Miller 2017) proved to be involved in calcite precipitation.

Altermann et al. (2006) estimated that cyanobacteria have been the main contributors to the production of carbonate rocks during almost 70% of Earth history. Since then, the importance of cyanobacteria in carbonation has been extensively investigated, with particular regards to the molecular mechanisms driving calcite precipitation in *Synechocystis* spp and *Synecococchus* spp (Lamérand et al., 2022; Görgen et al., 2021). In the course of the years *Bacillus subtilis, Pseudomonas nitroreducens* and *Bacillus licheniformis, Bacillus megaterium, Sporosarcina pasteurii*) (Ferral-Pérez et al., 2020; Rajasekar et al., 2021, Jarwar et al. 2022) became the most studied bacteria, and widely used in practical applications as soil consolidation and concreate healing, due to their ability to produce high amount of calcite within a short period of time (Bang et al. 2001; Dhami et al., 2013; Wei et al., 2015; Rajasekar et al., 2021).

Since 2000, the relevance of MICP in the wider frameworks of environmental microbiology and engineering applications has greatly increased. Some significant studies involving microbial induced calcite precipitation, applied to the geotechnical improvement of soil and concrete healing, have been published within the first decade of 2000s (Riding 2000; Visscher et al., 2000; Ben et al., 2004; Dupraz et al., 2004; Wright and Wacey 2005; Baumgartner et al., 2006). The interest of geotechnical potential use of MICP is growing today and it is fueling researches in the field of soil consolidation and use of microorganism in concrete healing. Bu et al. (2022) reviewed MICP soil improvement discussing in detail the mineralization mechanism, its technological importance and its effects on soil with different characteristics. As the concrete healing process through MICP is strongly dependent from the technological characteristics of the microorganism involved in the process, today many researches are focused on finding microbial strains with highly calcite precipitating properties. Of particular interest is the work of Sohail et al. (2022), who investigated the self-healing process in concrete through microbial induced calcite precipitation (MICP) carried out by a strain of *Bacillus cereus* isolated from soil in Qatar. However, soil stabilization as well as biocementation are not the only topics boosting the interest on MICP. Such an interest spans from the role of MICP in fossils formation and in the preservation of traces of life in ancient rocks, to the contribution of microorganisms to ancient and modern geochemical cycles of C and Ca, to the investigation on new methods to capture and store carbon, to the removal of the pollutant heavy metals or radionuclides (e.g. Ra, Sr) from environmental samples through co-precipitation with CaCO_3_ and to the use of carbonate biominerals to enhance oil recovery (Görgen et al., 2020).

The conspicuous number of research and review articles on MICP and its practical applications makes necessary an effort devoted to organize, summarize and quantify the recent development in the field in order to have a clear overall picture of the present state and future directions of the research. Bibliometric analysis, defined as the use of mathematical methods to analyze published articles in terms of quantity, quality and impact is an important and common method used to assess research activity on a certain topic (Sweileh et al., 2016). Further, it has proved to be a useful tool to quantitatively assess trends and patterns of scientific literature (Otte and Rousseau, 2002; Xiao et al., 2022). Bibliometric analysis can help to understand the research trend and development status by assessing the authors, institutions, countries and link strength based on keywords associated to publications and helps to pave a pathway for future studies (Li et al., 2017; Xiao et al., 2022).

Even though bibliometric analysis has been widely used in the field of environmental sciences, very few studies focus on environmental microbiology or geomicrobiology. Some examples of bibliometric analysis in the field of microbial ecology are given by Qi et al. (2019), who analyzed the research status and development trend of algae-bacteria symbiotic wastewater treatment technology between 1998 and 2017, and of Vanzetto and Thome (2019), who investigated the toxicology of nanoscale zero valent iron used in soil remediation by bibliometric analysis. However, to best of our knowledge, no studies have attempted yet to analyze the structure, links and future research avenues on MICP. This paper aims at exploring scientific literature on MICP based on keywords analysis, most productive authors, most cited published paper, most productive institutes and countries, collaboration between authors, countries and institutes, as well as to highlight the scientific journals interest for this topic over the past 22 years.

## 2. Materials and Methods

### 2.1. Data Source

Scientific output data was extracted from the Web of Science Core Collection (WoSCC), one of the most widely used daily updated databases in academic and bibliometric studies, allowing the download of full citation records (AlRyalat et al. 2019). WoSCC can provide comprehensive information for bibliometric analysis, including the Science Citation Index-Expanded (SCI-E) and the Social Sciences Citation Index (SSCI). In this study, WoSCC database was used to retrieve the related research on microbial-induced calcite precipitation (MICP) for the period from 2000 to 2022. The language of the analyzed papers is limited to English with an overall number of 1580 publications. We downloaded and exported the collected data (which include full records) in text format for further analysis. All data were exported within on August 30, 2022, to avoid deviations caused by the daily updates of this database.

### 2.2 Bibliometric network analysis

We performed the study of the global scientific research and trends on MICP through a bibliometric network analysis carried out according to Reuters (2008). We based our study on the visualized analysis of Mapping Knowledge Domain (MDK) and on the implementation of a Social Network Analysis (SNA) (Zou et al., (2018). While MKD can be used to establish a reference information and research basis for the application and development of methods of a chosen domain, SNA is defined as the process of investigating social structures using networks and graph theory (Otte and Rousseau, 2002; Zou et al., 2018). SNA and maps based on network data allow the application of systems thinking in bibliometric science. The outputs of such analysis are network maps and statistics based on the relationships among countries, journals, organizations, authors, and keywords related to the investigated topic (Chen et al., 2016).

In this study, we used the VOSviewer software (version 1.6.18) to perform the bibliometric analysis. This software, using the VOS (Visualization Of Similarities) mapping technique, is especially useful for displaying large bibliometric maps in an easy-to-interpret way while allowing the creation and exploration of maps based on bibliometric network data. The output results are displayed in clusters to visualize the existing connections among the analyzed bibliometric data. VOSviewer software is a useful tool for the elaboration of distance-based maps in which the distance between two items reflects the strength of the relation between the items. Unlike graph-based maps, in distancebased maps a smaller distance generally indicates a stronger relation (Van Eck, N.J., Waltman, 2010). Table 1 summarizes the main technical terms used by the software. We implemented a co-authorship, co-occurrence, and citation analyses were conducted to create distance-based network maps showing: (1) the co-authorship among researchers and collaborations of countries, (2) the co-occurrence of keywords, and (3) cited scientific journals.

**Table 1** Terminology used by VOS viewer software (Van Eck and Waltman, 2022).

## 3. Results and Discussion

### 3.1 Document types

As of 30^th^ August 2022, 1580 articles related to MICP was identified and classified into eight document types indexed in the Web of Science (Table 2). Article was the most frequently used document type, consist for 86.01% of the total publications, followed by conference proceeding papers (11.13%). Review papers contributed (4.11%), early access (1.9%), book chapters (0.06%), editorial materials (0.31%), corrections (0.12%), and meeting abstracts (0.06%), which showed lower numbers than article and conference proceedings. It is thus obvious that the article and conference proceedings with the largest proportion may greatly reflect the development trends and change in this research field. Some research reports on original works were classified as article and proceedings at the same time, which cause the sum of percentile in all types is greater than 100%.

**Table 2.** Document types of publications on Microbial induced calcite precipitation (MICP) from 2000 to August 2022.

### 3.2 Publications and citations trend analysis

The web of science database search resulted in 1580 publications on “MICP” in the period from 2000 to August 31^st^, 2022 (date in which the analysis was performed). Both the cumulative citation index and number of publications show an increasing trend in the investigated timeframe. Over the last 10 years, the number of publications has increased significantly, and the topic of calcite precipitation had been studied more frequently. From the year 2000 to 2015, as it can be seen in Figure 1, the number of publications dramatically increased and so does the citations. In 2010, only 10 papers were published on the topic of MICP but, in 2021, 333 papers were published worldwide covering this topic proving the emerging importance of the topic

**Fig 1.**
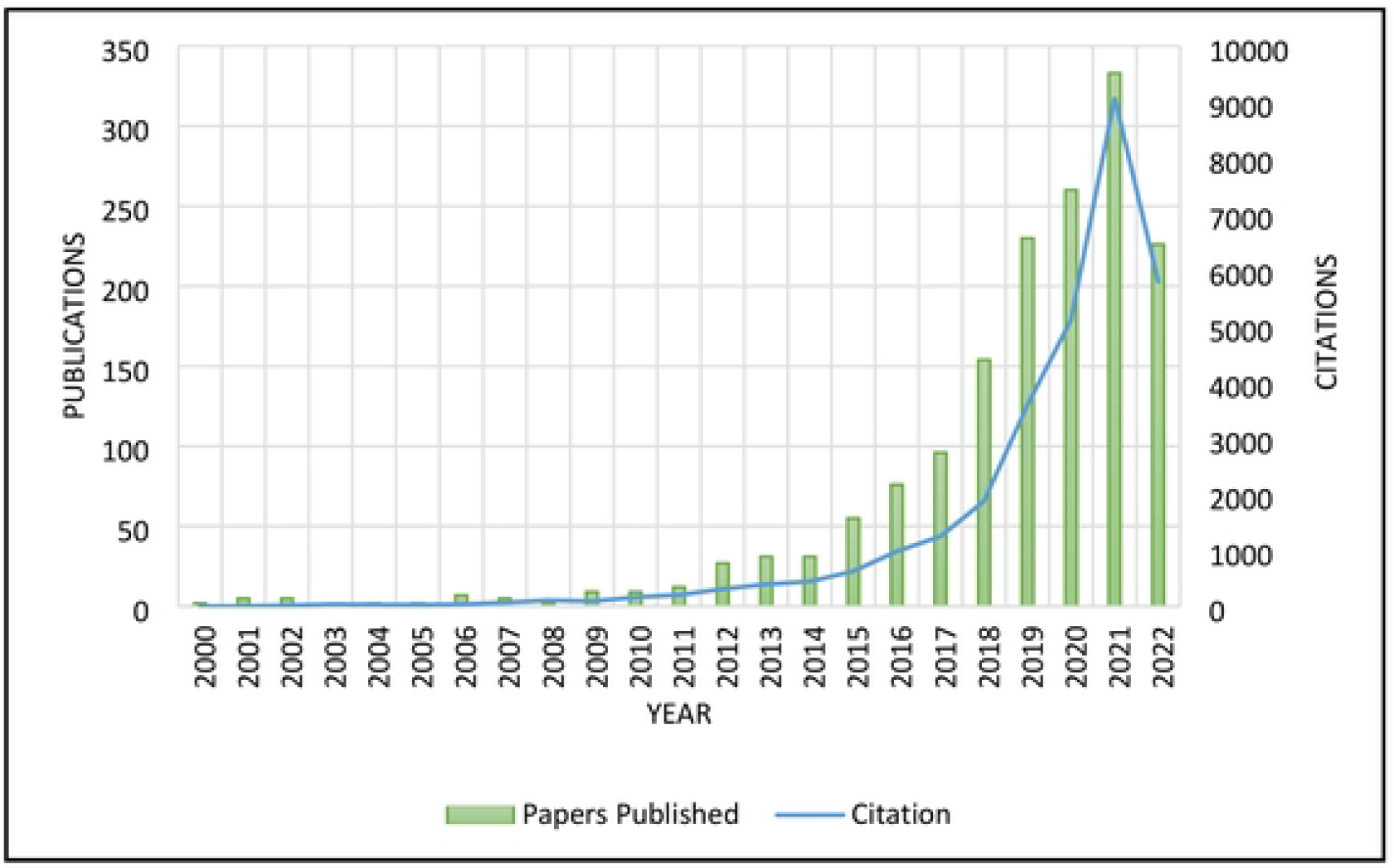
Number of publications and citations in the field of MICP from 2000 to August 2022.

### 3.3 Countries and collaborations network analysis

In 2000, only two papers were published on the topic of MICP and both were published by researchers from China. Meanwhile, 71 countries carried out research on MICP by the end of 2021, which indicates the increasing amount of research possibilities and interest this topic possesses. Among all participated countries, 9 countries have published more than 50 papers until August 2022 as shown in Figure 2. China is leading in research on MICP with total 648 publications, followed by United States (415), Australia (103) and India (87) respectively. The high output of publications in these countries can be explained by their strong economic strength and the huge investment in research, development, and innovation.

**Fig 2.**
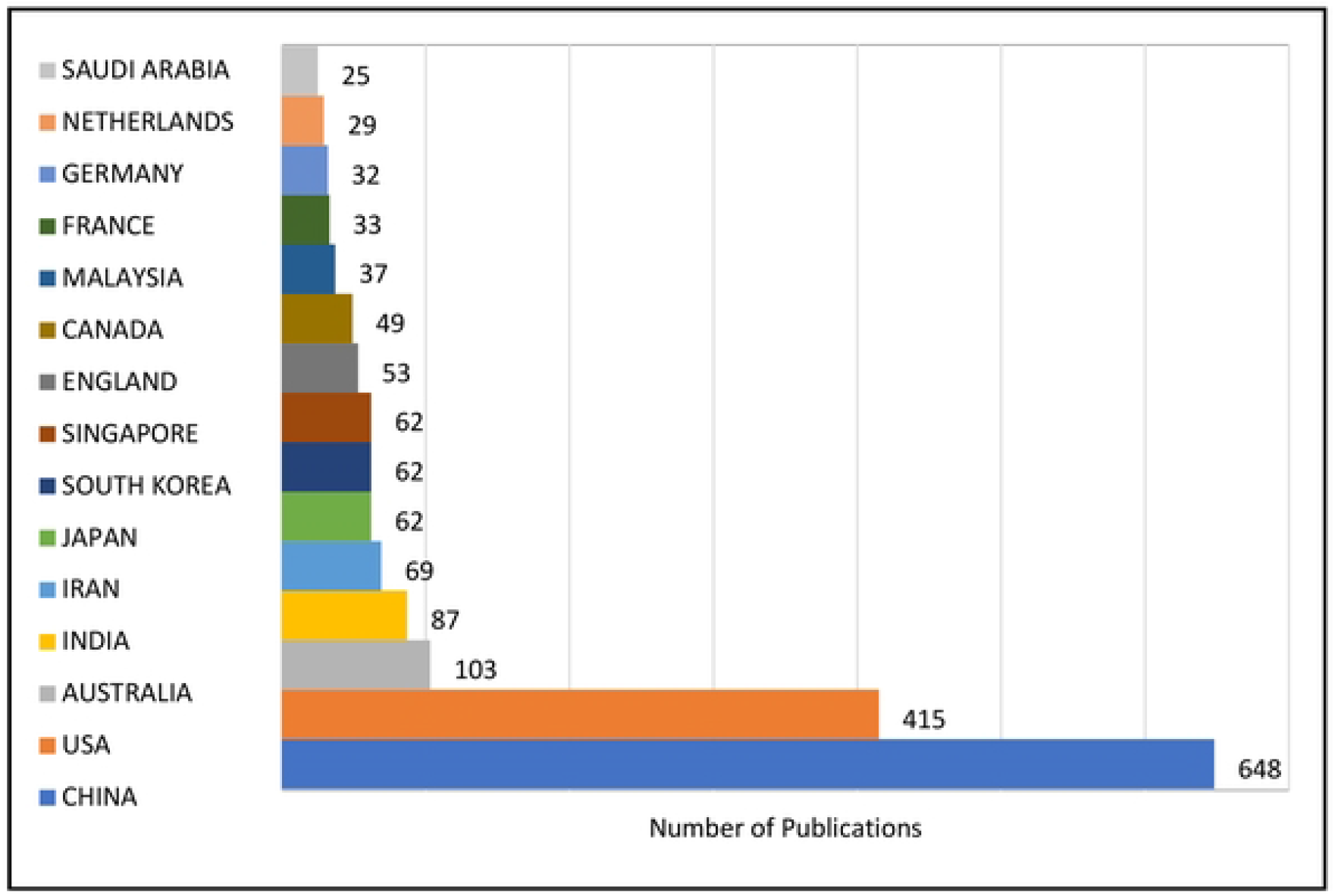
Top 15 countries with highest number of publications from 2000 to August 2022.

Academic cooperation between different countries or research institutions plays a guiding role in promoting dissemination of knowledge and academic exchange among scholars (Chen et al. 2020). The academic cooperation relationship among the top 10 countries of publications from 2000 to 2022 is shown in Figure 3. The nodes in the figure represent different countries, the lines connecting the nodes indicate international cooperation between countries, and thickness of the line represents the closeness of cooperation.

**Fig 3.**
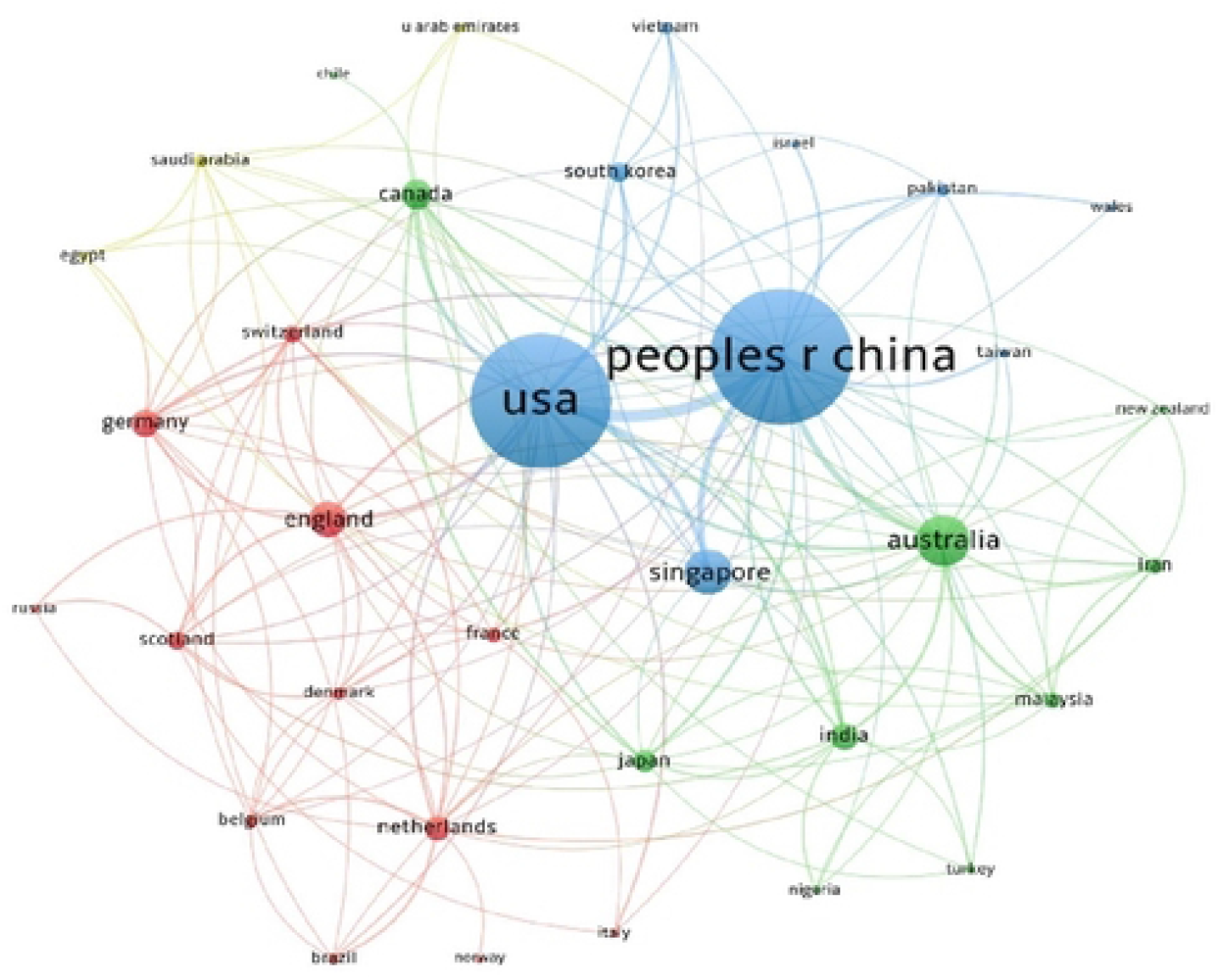
The academic collaboration networks among the top productive countries.

China and USA are the most active countries in global cooperation on research of microbial induced calcite precipitation and both are located at the center of the collaborative network. USA and China both cooperated with 27 countries to publish in MICP research, Australia ranked Second as cooperating with 17 countries followed by Netherlands (15), England (15), India (12) and Iran (10). China and USA had the closest collaborations and both countries collaborated very closely with Singapore. China also has cooperated closely with Australia. There is still much gap and loophole for the development of the cooperation between countries. Like, the USA and China, the two major countries with the largest publications, have never collaborated with Russia and Brazil.

### 3.4 Institutions and collaborations network analysis

More than 1270 institutions have contributed in researches on microbial induced calcite precipitation (MICP). The top 10 institutes accounting for 37.24% of total publications with 549 published papers. China has 5 institutes in top 10, followed by USA with 2 institutes. However, Japan, Singapore, Australia have 1 institute each in top 10. China university of Petroleum ranked first with 98 publications, followed by Nanyang technological University with 62 publications, Chinese Academy of Sciences with 59 publications and China University of Geoscience with 56 publications.

China University of Petroleum have more paper published in the field of MICP, but Nanyang Technological university Singapore have collaborated with 18 institutes worldwide (Figure 4), followed by Chongqing university China with 14 institutes, Jiangsu University China with 13 institutes, Chinese Academy of Sciences and University of Hawaii Manoa with 12 institutes. Most of the institutional collaborations are domestic or with local institutes. Cooperation between institutions from different countries is crucial to the development and dissemination of knowledge, thus international cooperation in this research fields needs to be improved.

**Fig 4.**
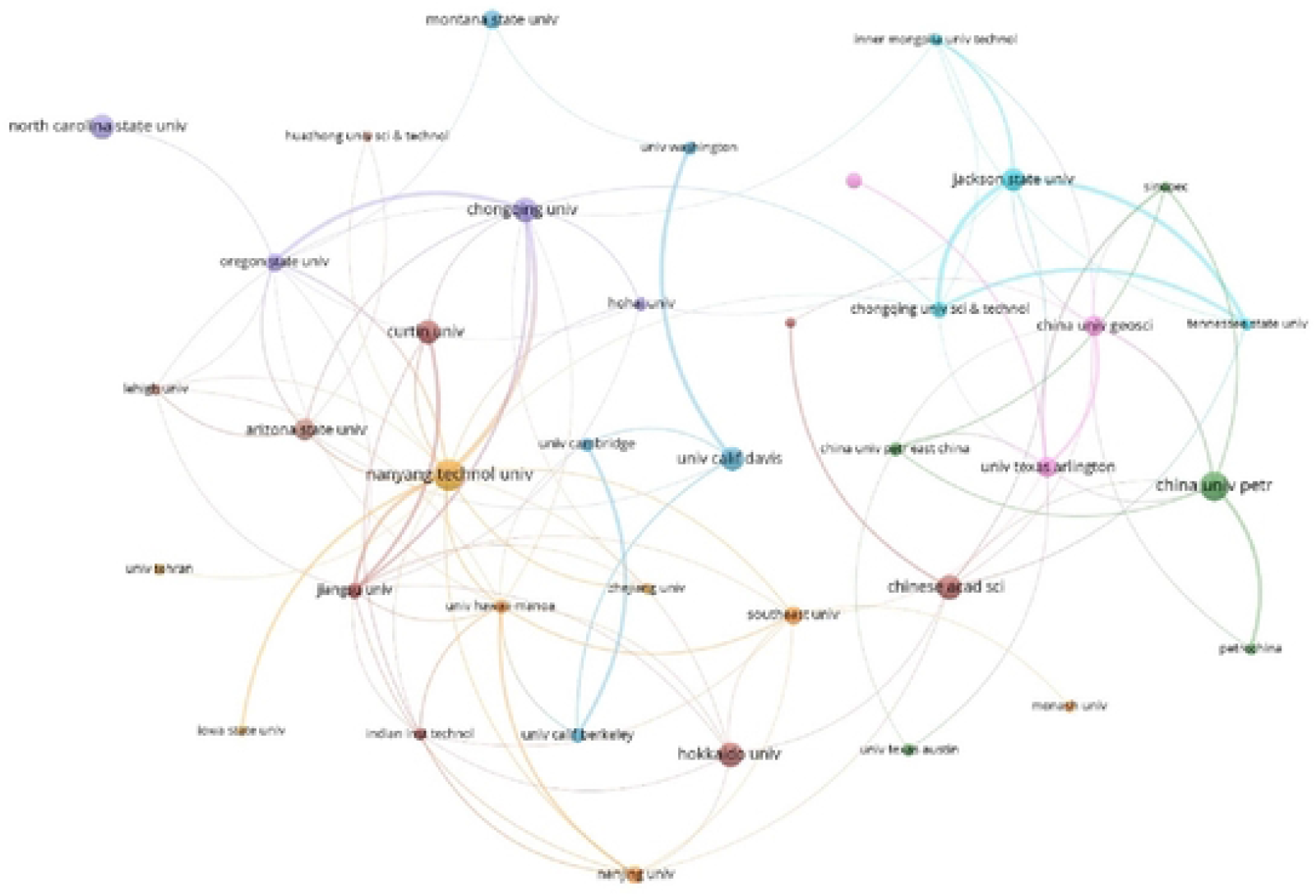
The academic collaboration networks among the top productive institutes.

### 3.5 Authors and collaborations network analysis

Almost 3900 authors have contributed to the research of microbial induced calcite precipitation (MICP) from 2000 to 2022. In China, researchers have produced more researches in this field (Figure 5), as 4 out of 10 top productive authors are from China or associated with Chinese institutes, whereas 3 researchers are from USA, 2 from Japan and 1 from Singapore. Chu Jian from Nanyang Technological University Singapore leads with 51 publications, followed by Kawasaki Satoru from Hokkaido University Japan with 40 publications, Montoya Brina from North Carolina State University USA and Cheng Liang from Jiangsu University, both ranked third with 36 Publications.

**Fig 5.**
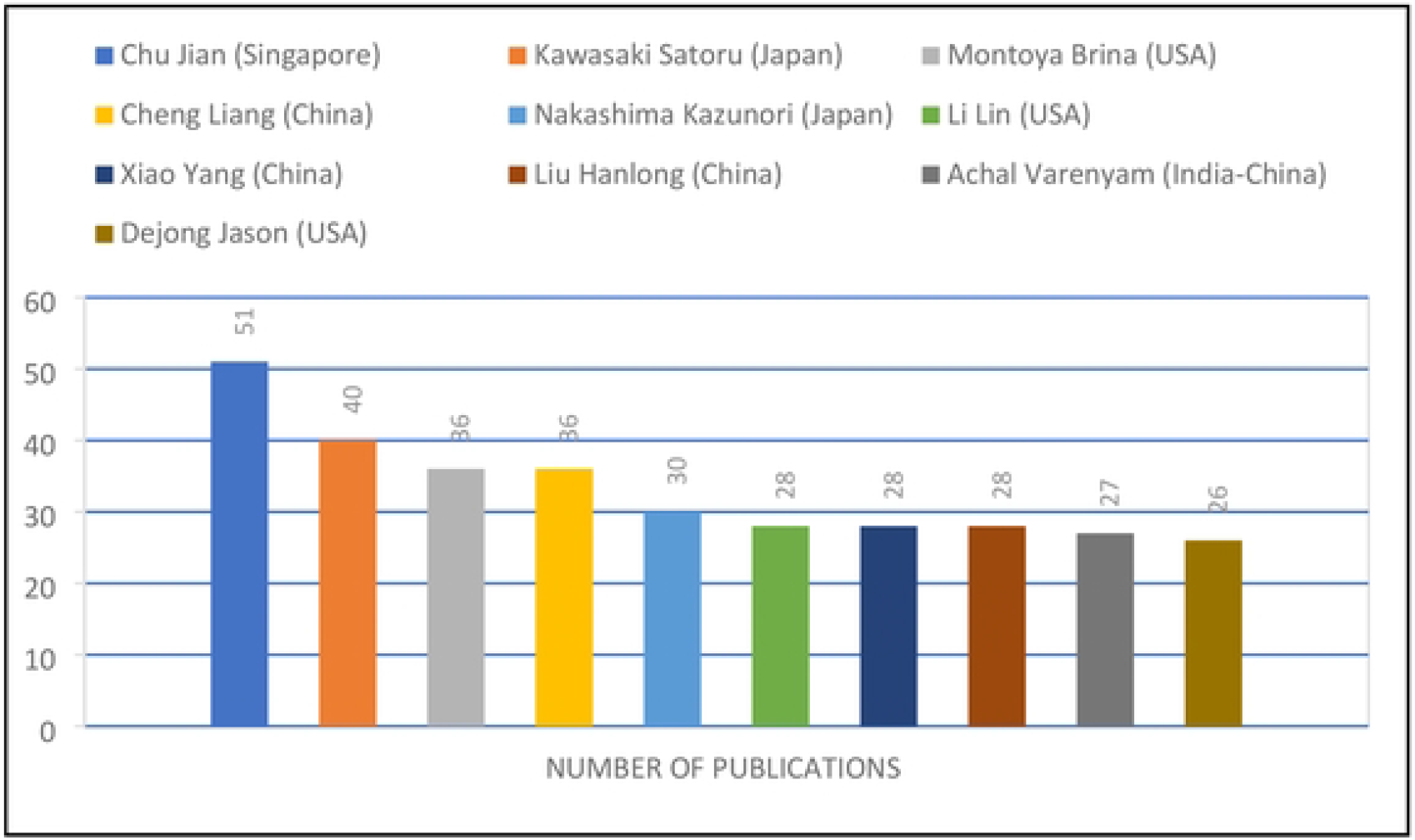
Most productive authors ranked by number of publications.

Figure 6 shows the collaboration between top productive researchers. This cooperation network includes a large cooperation network and several small cooperation networks, which are not connected with each other. Chu Jian from Nanyang Technological University of Singapore collaborate with 23 different researchers around the world and tend to be most productive and collaborative author. Liu Han Long from Chongqing University China and Cheng Liang from Jiangsu University China were second most collaborative researchers in the field of MICP and have collaborated with 16 researchers each. Xiao Yang from Chongqing University China collaborated with 15 researchers. Surprisingly, Kawasaki Satoru (Hokkaido University, Japan), the second most proliferous author in this field after Chu Jian (Nanyang Technological University, Singapore) had collaborated with only 8 researchers.

**Fig 6.**
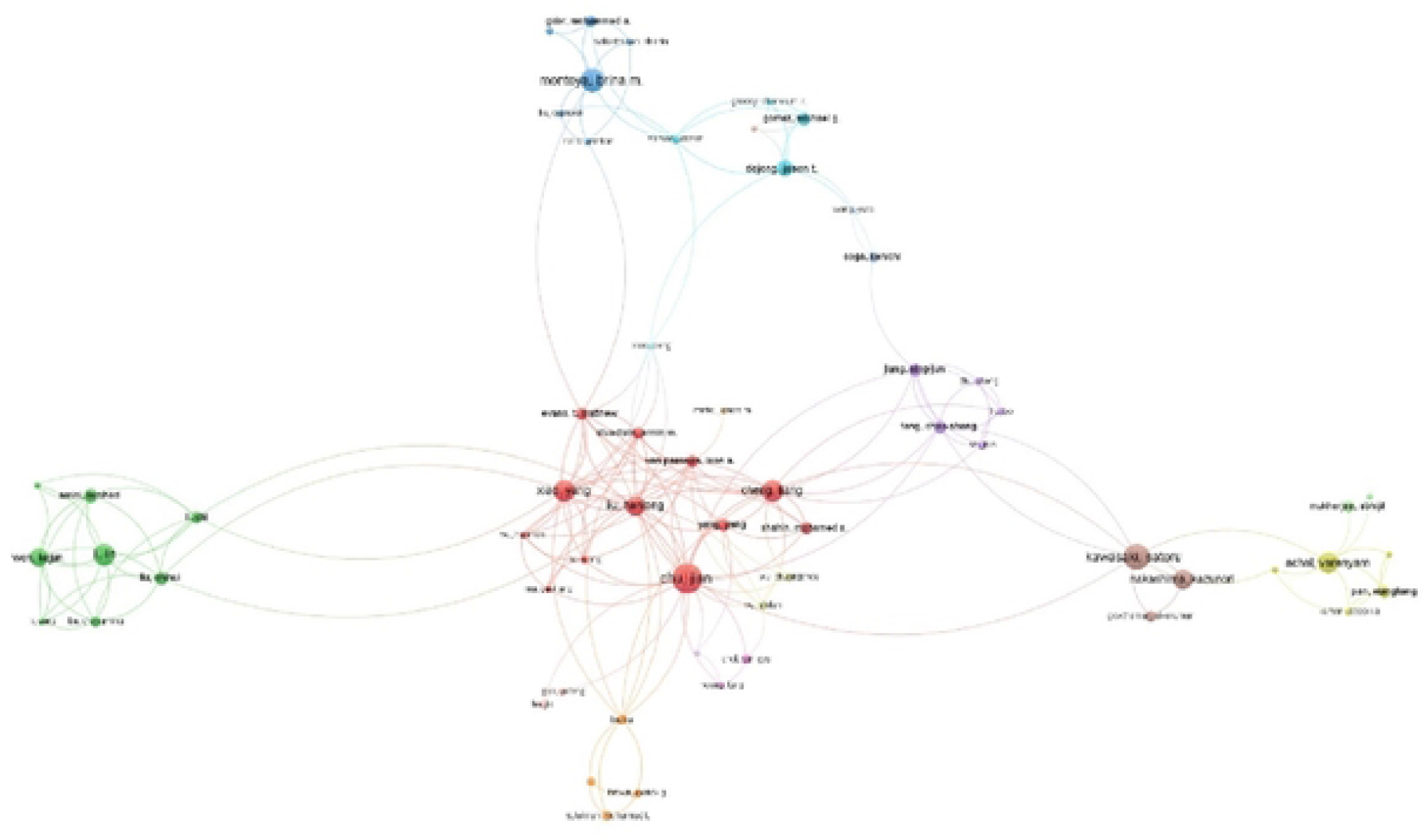
The academic collaboration networks among the top productive authors.

### 3.5 Most cited articles

Citation analysis is an important indicator to measure the quality of publications, and it represents both the interest raised by a research topic within academic circles and the attention scientific community pays to the work of a researcher (He et al. 2019). The top 10 most cited papers from 2000 until date in the field of MICP are listed in Table 3, where the papers are classified according to the title, type, first author’s name, journal, total citations and year of publication.

**Table 3.** Top 10 most frequently cited publications from 2000 to 2022

Our analysis highlight how the top cited papers were published by journals in the geotechnical engineering area (Geotechnical and geoenvironmental engg.) Just one paper was published by a journal of the biological area (Applied microbiology). This result shows that, despite the fact that MICP is a geo-biological process, practical applications in soil improvement as well as construction materials reparation obtain the highest interest from the scientific community.

### 3.6 Journals network analysis

Figure 7 shows the top journals, in term of numbers of publications on MICP. A total of 521 journals, with Impact Factor (IF) ranging from 2 to 7, published papers on this topic. *Construction and building materials* (IF: 6.141) is the most productive journal, as published 65 papers in this research area, accounting for the 4.12% of total papers published on this topic. It is followed by *Petroleum science and engineering journal* (IF:4.346) with 61 published articles, accounting for 3.86% of total publications. *Journal of Geotechnical and Geoenvironmental Engineering* published 51 papers, followed by *Geotechnical special publication*, a conference proceeding by American Society of Civil Engineers (ASCE) library, published 47 papers. These results confirm that the engineering area is the field actually leading the research about MICP.

**Fig 7.**
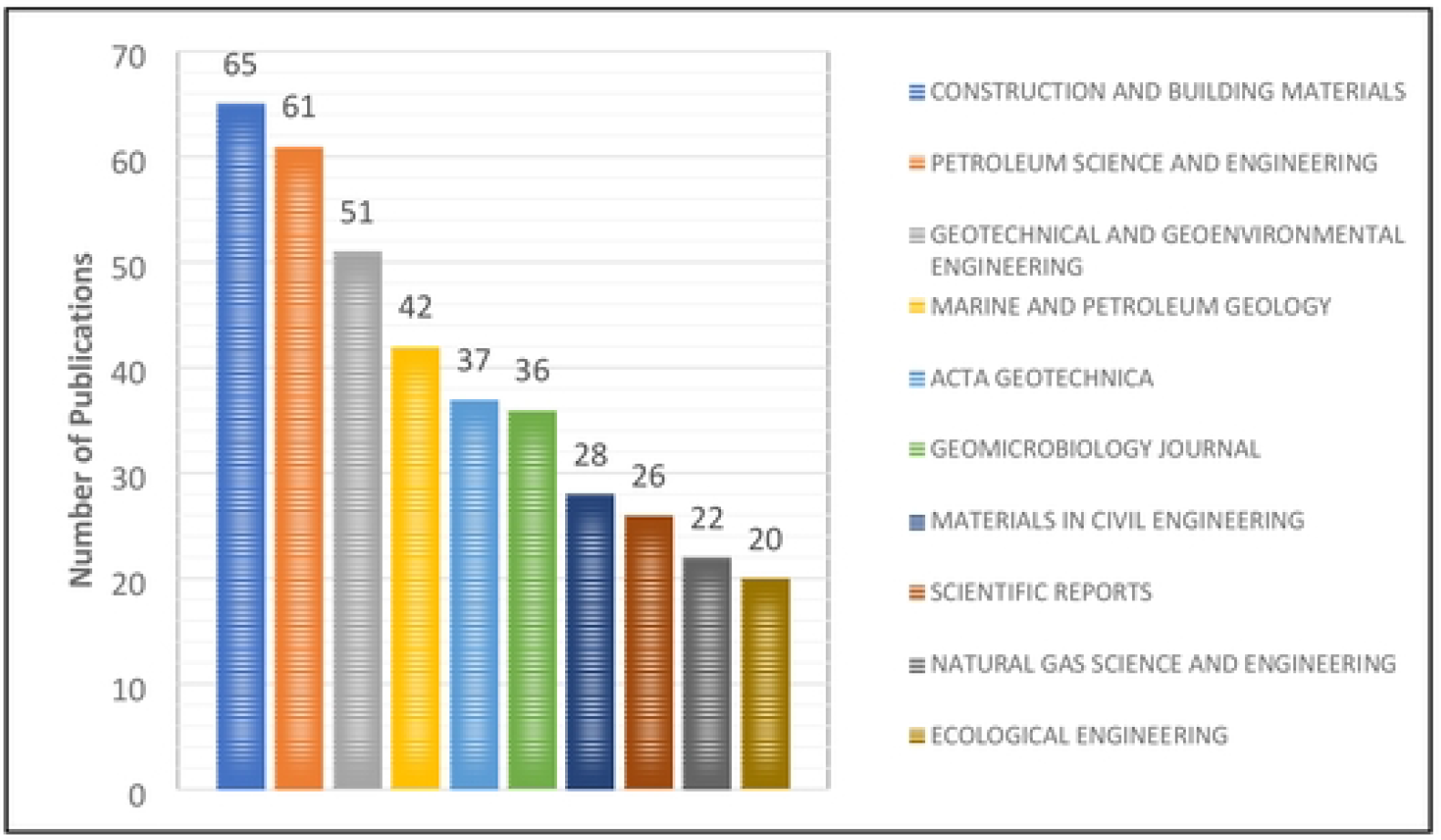
Top journals publishing microbial induced calcite precipitation (MICP).

### 3.7 High frequency keyword analysis

The research hotspots and overall trends in the field are accurately revealed by the analysis of the high frequency keywords, their links, total link strength, (Qi et al. 2019). The analysis of the keywords related to MICP generated 3443 results. Among them, only 299 met the threshold of at least five co-occurrences, which is 8.68% for total count. 143 keywords appeared 10 times and 522 keywords appeared 3 times which is 4.15% and 15.16% respectively. MICP used 251 times with total link strength of 1600 as shown in Table 4, followed by carbonate precipitation, which co-occur 213 times with total link strength of 1361.

**Table 4.** Top 10 authors keywords, frequencies and total link strength.

The co-occurrence network map of keywords is shown in Figure 8, in which the bigger is the size of the circle, the higher the co-occurrence of an item. Moreover, the shorter is the distance among items, the stronger their relation. MICP linked with 257 researched items, followed by carbonate precipitation, cementation, improvement, sand etc.

**Fig 8.**
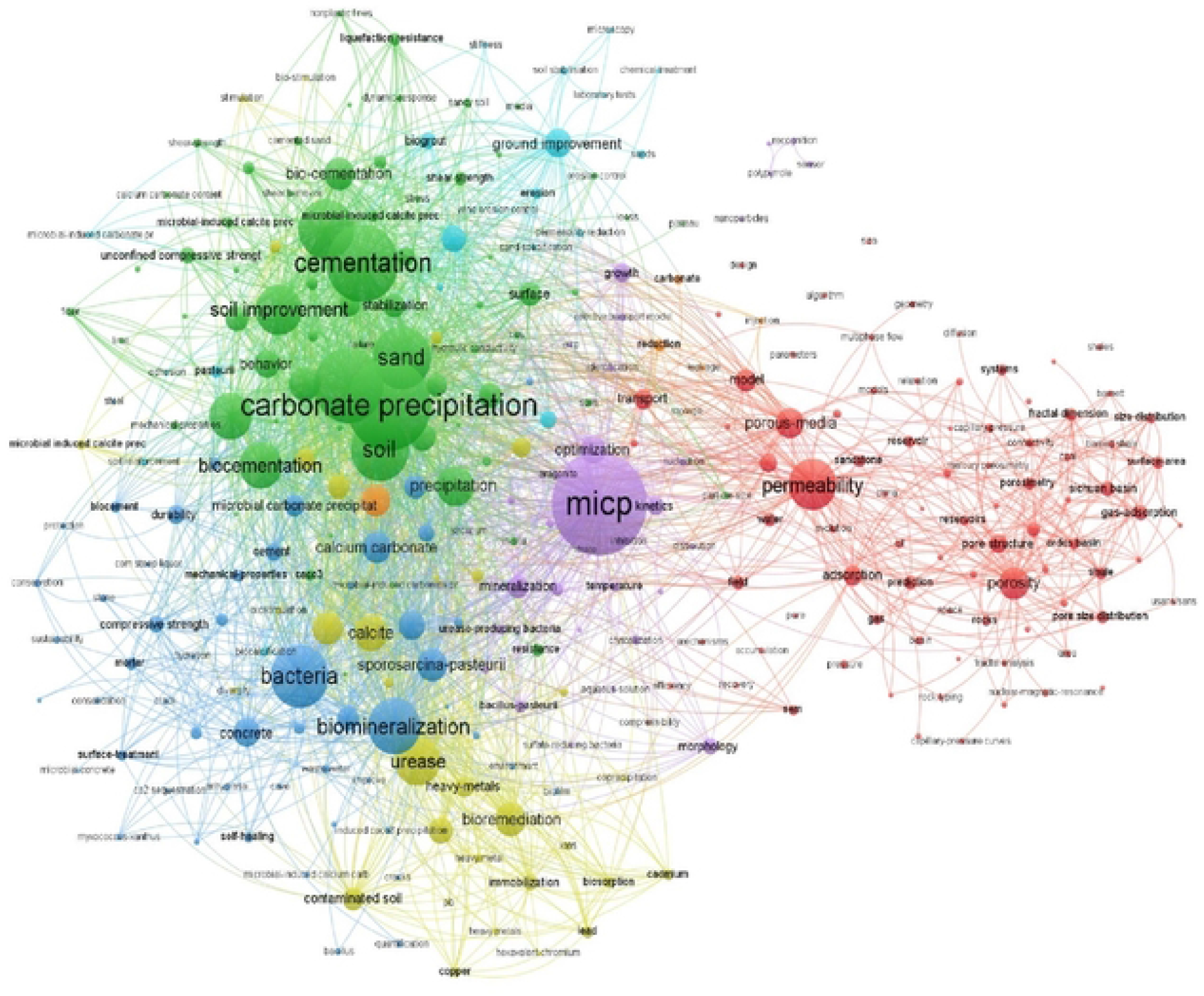
Keyword co-occurrence network map based on total link strength.

## 4. Conclusions

In this study we explored different aspects of research on microbial induced calcite precipitation (MICP). We performed a bibliometric analysis based on 1580 publications, from the year 2000 to 2022, retrieved from web of science core collection database. The bibliometric analysis allowed the quantification and visualization of the multi-disciplinary development of MICP, the understanding of the most productive authors, institutes, countries, journals and the future research directions. The most active research field is geotechnical engineering and China is the leading country. Russia and Brazil are not highly involved in international networks, showing the presence of a great gap for the development of cooperation in this research field. An extension of the research network is needed to foster the progress in MICP field.

Concrete healing or bio cementation, soil improvement and consolidation through MICP are the topics of focus. Further, our results show that not always a high scientific production by an author occurs in presence of a wide network of international collaboration.

We believe our paper takes important information to the scientific community, as, for the first time, relatively objective and comprehensive bibliometric analysis on the research field of MICP has been carried out. Just like other studies, there are still some limitations due to incomplete search item and some relevant articles may have been missed. In addition, only articles written in English were included, so there is a chance that some important articles, written in other languages, were missed too.

## Data Availability Statement

Not applicable.

